# Electrospinning of hydroxypropyl chitosan nanofibers for bone regeneration application

**DOI:** 10.1101/2025.01.05.631378

**Authors:** Junmei Wang, Jie Xiong, Guozhao Li, Ziliang Zhou, Yanyan Yang, Lu He

## Abstract

To study the effect of electrospinning nanofiber mats synthesized using hydroxypropyl chitosan in conjugation with poly (vinyl alcohol) on the osteogenesis in MC3T3 cell, we fabricated different ratios of hydroxypropyl chitosan (HPCH) and poly (vinyl alcohol) to electrospin the nanofiber mats, the mechanical and physical properties of nanofiber mats were tested by using SEM, FT-IR and conventional methods. Then the effect of the electrospinning mats on osteogenesis in MC3T3 cell were evaluated by Alizarin Red stain, ALP stain and expression of relevant genes. We found that the nanofiber mats fabricated by hydroxypropyl chitosan (50%) and poly (vinyl alcohol)(50%) can induce osteoblast differentiation apparently in MC3T3 cell.

## INTRODUCTION

Bone defection is a common problem in dentistry coming from periodontal disease, teeth extraction and trauma, then how to prevent the resorption of bone and preserve the alveolar ridge are crucial.

Restoring sufficient bone volume to enable ideal bone regeneration on deficient sites originated from osteogenesis, osteoinduction and osteoconduction. Based on this, a variety of biomaterials are used to stimulate bone regeneration from these three aspects.

Electrospun nanofiber membranes have advantages of specific surface and mimicking extracellular matrix (ECM), then be used widely in nanofiber biomaterials. (Tsai et.al.,2015;Ray et al.,2017). Some researchers developed three types of cytocompatible nanofiber membranes using of chitosan/gelatin/PEO (CGP) by electrospinning to enhance the cell proliferation and infiltration(Kuo et al.,2018).There’s evidence demonstrate that Electrospun nanofiber membranes fabricated by hydroxypropyl cellulose (H), chitosan (C) and polyethylene oxide (P) possess properties of nontoxic, anti-fibroblast adhesion and anti-bacterial(Che et al.,2020)

Hydroxypropyl chitosan (HPCH) as a semisynthetic product from etherification and propylene oxide of chitosan under alkali conditions is a desirable choice for tissue regeneration, because of containing the primary amine group at C2 position thereby the properties of chitosan were also maintained. After the chemical modification, HPCH exhibits good hydrophilic abilities which make it a desirable scaffold material for tissue regeneration compared to chitosan. Because of excellent film forming, antimicrobial, biodegradability ability, HPCH has been applicated broadly in biopharmaceuticals fields. However, the ability of bone regeneration hasn’t been proved deeply (Chimenti et al.,2013; Ghannoum et al.,2015; Piraccini et al.,2018).

Poly (vinyl alcohol) (PVA) as a highly hydrophilic and biocompatible semicrystalline polymer has also been widely used in electrospinning to form excellent nanofibers. But its applications were tied because of lower strength and high water-solubility, so chemical or physical crosslinking of PVA can be applied to enhance the mechanical properties of nanofibers (Saad et al.,2015;Enayati et al.,2018;Xue et al.,2016;Alavarse et al.,2017). In this study, we explore the morphology and mechanical of nanofiber mats through physical crosslinking of PVA and HPCH, the nanofiber mats were used to induce osteoblast differention in MC3T3 cell.

Therefore, the aim of this study is to investigate the potential role of electrospinning nanofiber mats fabricated by hydroxypropyl chitosan in conjugation with poly (vinyl alcohol) on osteogenesis in MC3T3 cell.

## Material and methods

### 1. Material

Hydroxypropyl chitosan (HPCH) (average Mw 80k g/mole) and Poly (vinyl alcohol) (PVA) (average Mw 125k g/mol) was purchased from Sigma-Aldrich Company, USA. Dulbecco’s modified eagle’s medium (DMEM), fetal bovine serum (FBS), penicillin and streptomycin were purchased from Gibco (Rockville, MD, USA). 6-well and 96-well plates were obtained from Corning Life Science (St Lowel, MA, USA).

## 2. Method

### 2.1 Polymers Solutions Preparation

HPCH (20g) and PVA (20g) were dissolved in distilled water (100 mL) to make the solution respectively. The solutions were stirred with a magnetic stirrer mildly for 4h at 60°C,then were slowly cooled down to room temperature.

### 2.2 Blended Solutions of HPCH/PVA

The previous solutions were blended with a magnetic stirrer for 24h at 60°C to obtain different ratios of HPCH and PVA such as 100/0, 75/25, 50/50 and 0/100, Table I lists the different compositions and volumes of the blended solutions of HPCH/PVA.

**Table 1.** Composition and volume of blend solution of HPCH and.

| Sample name | Composition (%) |  | Volume of solution (ml) |  | Total volume |
| --- | --- | --- | --- | --- | --- |
|  | HPCH | PVA | HPCH | PVA |  |
| HPCH | 100 | 0 | 3 | 0 | 3 |
| HPVA1 | 75 | 25 | 2 | 1 | 3 |
| HPVA2 | 50 | 50 | 1.5 | 1.5 | 3 |
| PVA | 0 | 100 | 0 | 3 | 3 |

### 2.3 Electrospinning Process

Electrospinning of the solutions was performed at environmental temperature of 32 °C, the solution was placed in a 5ml syringe: high voltage (30 kV) was applied to initiate the process. The feeding rate was controlled by the syringe pump (0.1 mL/min) and the tip-to-collector distance (TCD) was adjusted to 10 cm. The cell slides were adjacently put on a flat collector to obtain the nanofiber, then the nanofiber-coated cell slides were kept for using in a cabinet drier at room temperature.

### 2.4 Electrospun Nanofibers mats

The morphological structure of electrospun nanofibers was monitored using scanning electron microscopy (SEM) (Amray Model 1820 Scanning Electron Microscope, UK). The bonding configurations of the samples were characterized by means of Fourier-transform infrared (FT-IR, TENSOR 27, Bruker, Germany). Mechanical properties were measured by applying tensile loads to specimens prepared from the electrospun nanofiber mats of HPCH/PVA (100/0, 75/25, 50/50, 0/100). Specimens of 25 cm^2^ area were used for mechanical measurements. Mechanical properties were also determined by a universal material testing machine (Tinius Olsen model H25 KT, England) at ambient temperature and a relative humidity of 65% with an elongation speed of 10 mm/min.

## 3. Cell culture

MC3T3 cells were purchased from the American type culture collection (Manassas, VA,USA) and were maintained in DMEM supplemented with 10% heat-inactivated FBS, penicillin(100units/mL) and streptomycin(100μ g·mL^−1^) at 37°C in a humidified atmosphere.

### 3.1 Alizarin Red Stain

MC3T3 cells were cultured in nanofiber-coated 24-well slides for 20 days. Cells were fixed in 4% paraformaldehyde (SIGMA−Aldrich) for 15 min at room temperature. Then were washed with phosphate-buffered saline and incubated with Alizarin Red (SIGMA−Aldrich) 40 mM for 30 min. Cells were rinsed five times with ultrapure water to eliminate unbonded stains. Images of each cell slide were taken, and calcium deposits with representative images were quantified using Image J software.

### 3.2 ALP Stain and ALP activity assay

ALP was conducted to assess the effect of nanofiber mats on MC3T3 osteogenic differentiation, MC3T3 were seeded in nanofiber-coated 24-well slides for 20 days at 2 × 10^4^ per well. Alkaline phosphatase staining was followed by the kit instruction. Cells with alkaline phosphatase activity were dyed purple-black or blue-black. The results were subsequently observed under phase-contrast microscope and the representative images were preserved. The ALP activity was quantified photometrically at 405 nm by ALP assay kit.

### 3.2 Reverse Transcription-PCR

MC3T3 were seeded in nanofiber-coated 6-well slides at a density of 2 ×10^4^cells/well. After 20 days, Total RNA was extracted from MC3T3 cell using Trizol reagent according to the instructions of the manufacturer. Reverse transcription was performed using corresponding kits. Real-time PCR was performed with SYBR Green detection using specific primers in technical replicates. The mRNA levels were normalized to GAPDH mRNA. Changes in target gene expression levels were calculated using the comparative ΔΔCT method. The primer used are showed in Tabel 2.

**Table 2.**
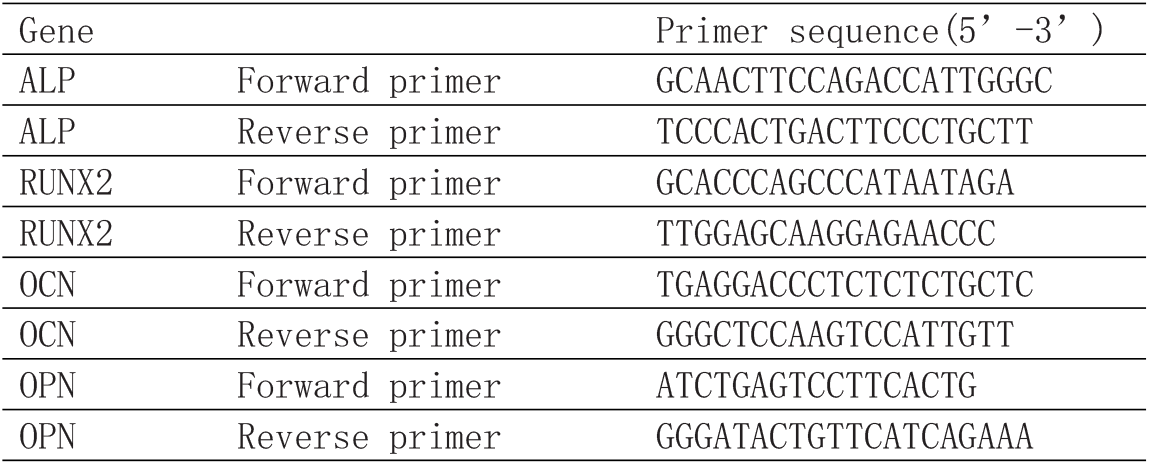
Nucleotide sequences of primers used for qRT-PCR.

### 3.4 Western blotting

The MC3T3 cells were seeded in nanofiber-coated 6-well slides at a density of 200 × 10^3^ cells per well. After 20 days the whole cell lysates were isolated using RIPA buffer and centrifuged at 12000 r/min under 4°C.The total protein were collected using Bradford method. After been denatured, SDS-PAGE electrophoresis (12% separating gel and 5% stacking gel) was used to separate proteins.

The obtained protein was transferred to nitrocellulose membranes and blocked in 5% nonfat milk for 1 h, then was incubated with the indicated primary antibody of RUNX2 at 4°C overnight for testing.

## 4 Results and discussion

### 4.1 Optical microscope (OM) and Scanning Electron Microscope (SEM)

The morphology of nanofiber mats fabricated by electrospinning were observed under Optical microscope (OM) and Scanning Electron Microscope (SEM), Figure1 showed the OM micrographs of the nanofiber mats produced from different ratios of HPCH and PVA. From the micrograph we can see that HPCH and HPVA1 fabricated bead-figured biomaterials on the cell slides, there’s no evidential nanofibers were found. HPVA2 and PVA can fabricate crosslinked nanofibers and present smooth surface. Under Scanning Electron Microscope (SEM), the diameter size and porosity were totally different fabricated by HPVA and PVA as shown in Figure2. The multi-plane of nanofibers from HPVA2 were captured under SEM in Figure3.

**Figure 1.**
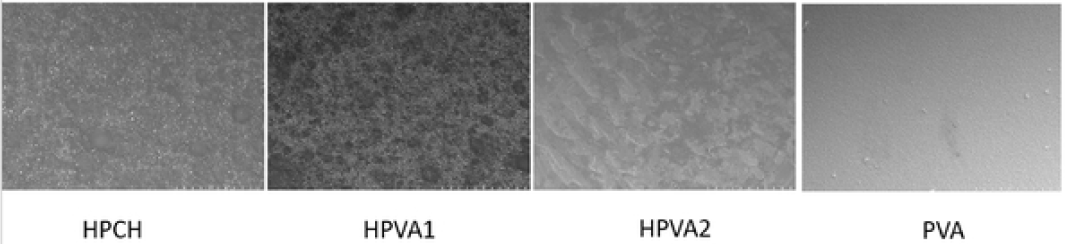
OM images of electro-spun mats with different ratio of HPCH and PVA(X200).

**Figure 2.**
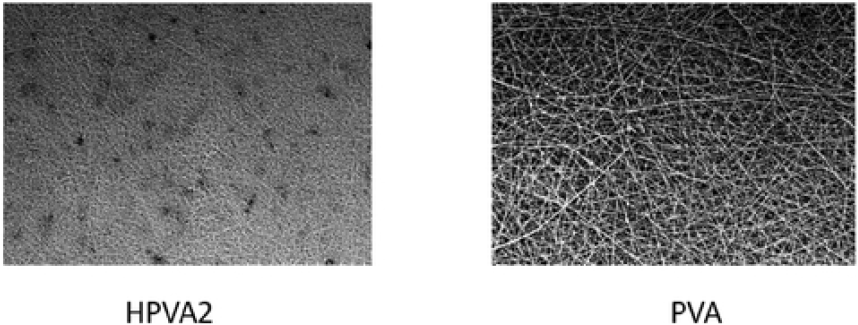
SEM images of electro-spun mats of HPVA3 and PVA4(X800).

**Figure 3.**
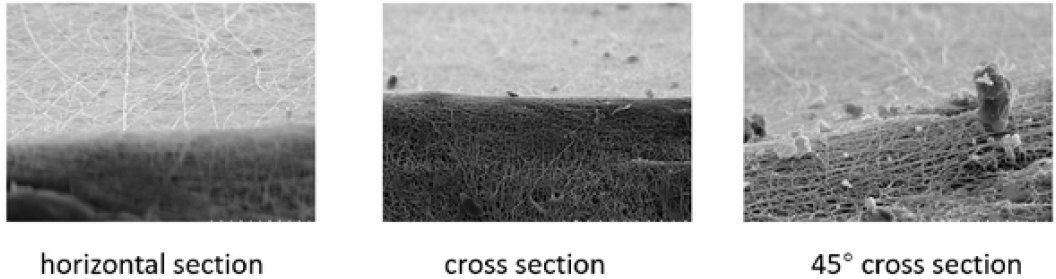
Different section of electros-pun mats with HPVA2 (X2000).

### 4.2 FT-IR Spectra of nanofiber Mats

Based on the upon results, HPCV2(50/50) simplified as HPCV was chosen for further study. Figure 4 illustrates the FT-IR Spectra results of PVA, HPCH and HPCV, FTIR spectra of nanofiber mats shows a peak at 3256-3259 cm^−1^ implying OH bonds and a peak at 2901–2920 cm^−1^ implying C–H bonds. There is not much FTIR spectra change among PVA, HPCH and HPCV. The results demonstrated that the blend solution didn’t change the original physical construction of HPCH and PVA, there’s no new configuration was produced in this procedure.

**Figure 4.**
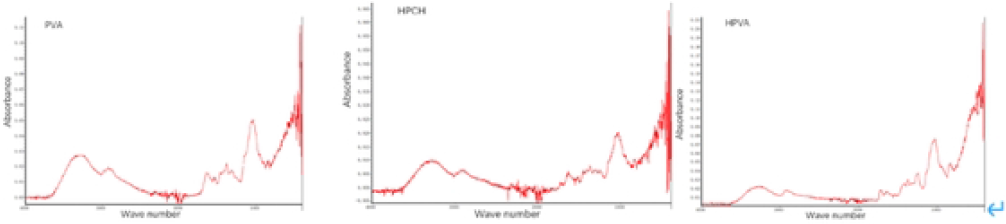
FT-IR spectra of PVA, HPCH, HPVA

### 4.3 Mechanical Properties of nanofiber mats

Tensile and stress forces were tested from nanofiber mats provided by HPCH and PVA. As shown in Table 3, the tensile and stress force were improved by blending the solution of HPCH and PVA. The ideal ratio is 50/50 which can provide proper mechanical properties.

**Table 3.**
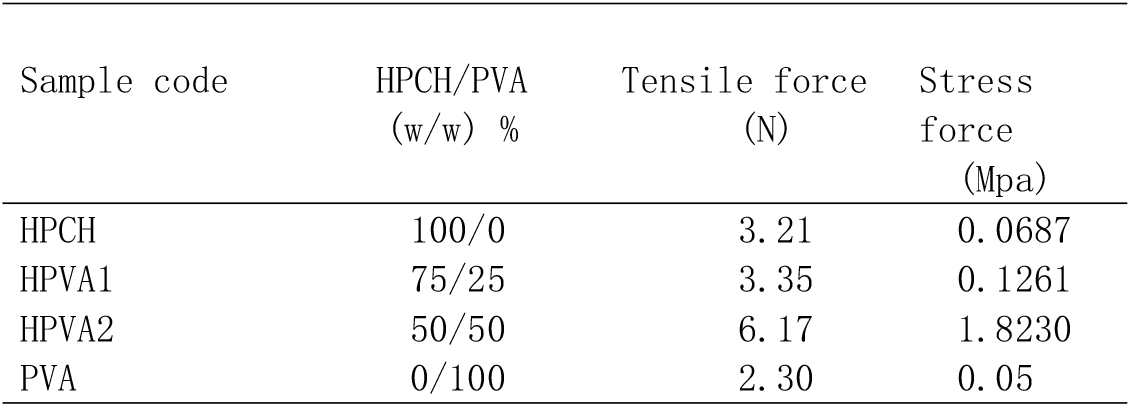
tensile and stress force of HPCH/PVA.

### 4.4 cell culture results

#### 4.4.1 Nanofiber Mats can induce osteogenesis

To evaluate the effect of nanofiber mats on osteogenesis in MC3T3 effectively, the cell-culture with nanofiber mats were divided with PVA, HPCH and HPVA (50/50). Then the nanofiber mats-coated cell slides were put in 6-well, 12-well and 24-well plates after 48-hour disinfection by ultraviolet light. As shown in figure 4, MC3T3 proliferated well around the nanofiber on the cell slide under the observation of Optical microscope (OM) and Scanning Electron Microscope (SEM).

Alizarin Red Stain and ALP staining showed that the nanofiber mats from HPVA can induce the osteogenesis of MC3T3 in Figure5 and Figure6.

**Figure 5.**
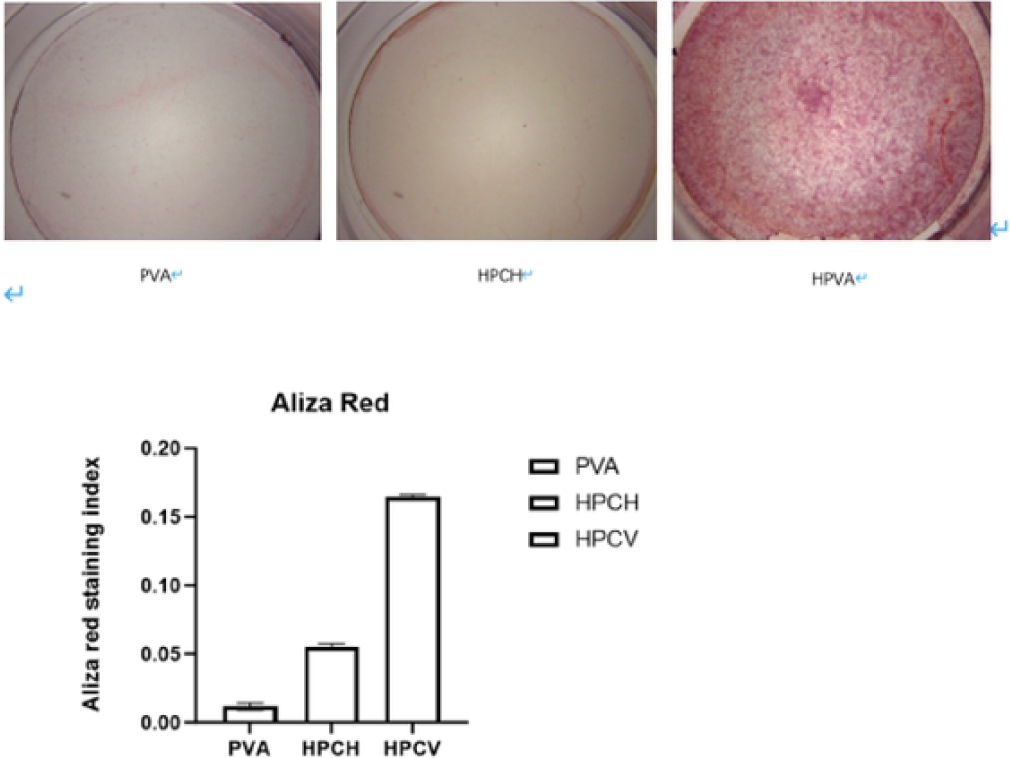
Results of alizarin staining(20 days)

**Figure 6.**
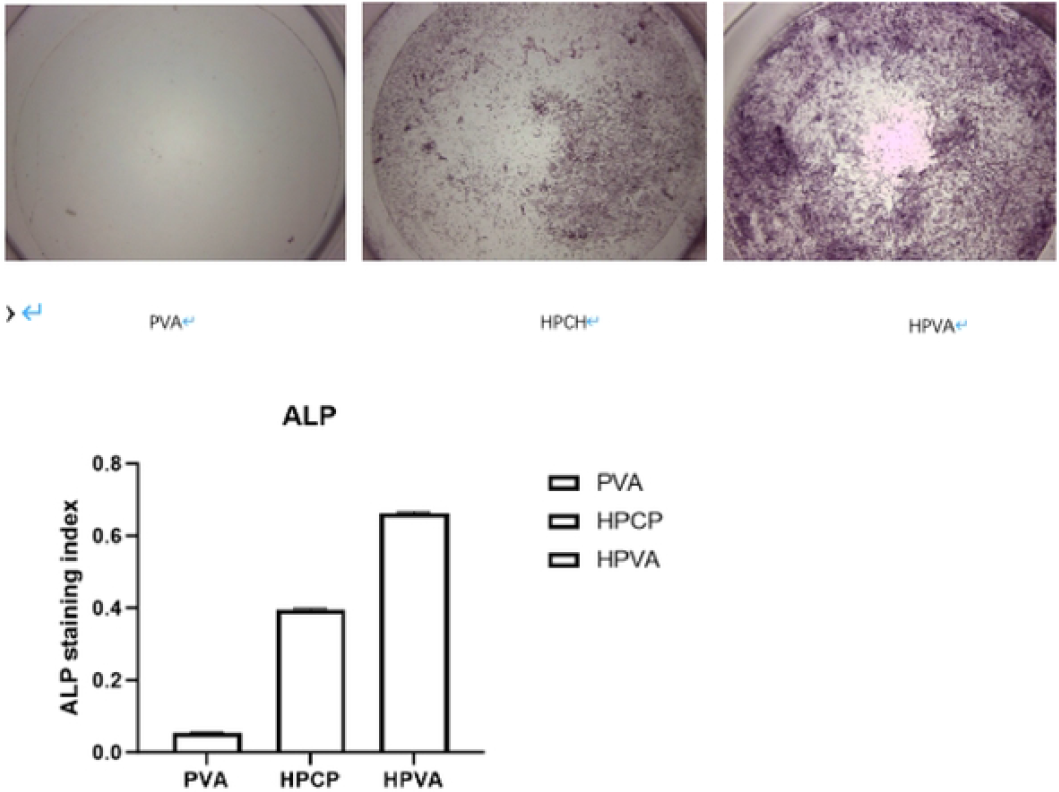
Results of ALP staininc (20 days)

**Figure 7.**
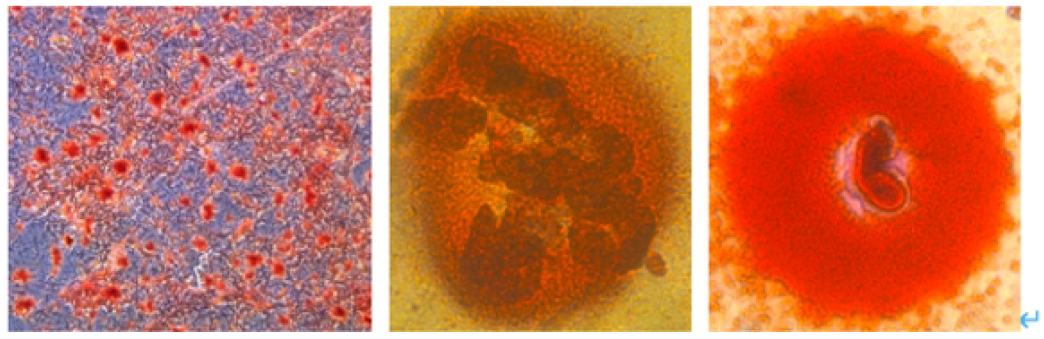
Red stained calcified node and osteoblast

#### 4.4.2 Nanofiber Mats induce osteoblast-associated gene expression

To further explore the effect of nanofiber Mats in osteoblast differentiation, we analyzed the relative expression level of osteoblast-associated genes. The expression level of osteoblast-associated genes was induced by the nanofiber mats (Table 4).

**Table 4.** the relative gene expression level.

| Group | Relative expression level |
| --- | --- |

|  | ALP | RUNX2 | OCN | OPN |
| --- | --- | --- | --- | --- |
| PVA | $1.000 \pm 0.000$ | $1.000 \pm 0.000$ | $1.000 \pm 0.000$ | $1.000 \pm 0.000$ |
| HPCH | $4.818 \pm 0.035^*$ | $6.692 \pm 0.010^*$ | $4.732 \pm 0.038^*$ | $5.374 \pm 0.025^*$ |
| HPVA | $5.423 \pm 0.023^\Delta$ | $8.282 \pm 0.005^\Delta$ | $6.386 \pm 0.012^\Delta$ | $7.064 \pm 0.008^\Delta$ |

\* $P < 0.05$ vs PVA group; $^\Delta P < 0.05$ vs HPCH group.

#### 4.4.3 Nanofiber Mats induce activation of RUNX2

Runt-related transcription factor 2 (RUNX2) is an essential transcription factor of osteoblast and trigger the expression of other osteoblast-related genes. As an important factor of osteoblast differentiation, the level of RUNX2 can indicate the osteogenesis of MC3T3. In Figure8 the transcriptional activity of RUNX2 increased apparently on HPVA nanofiber mats.

**Figure 8.**
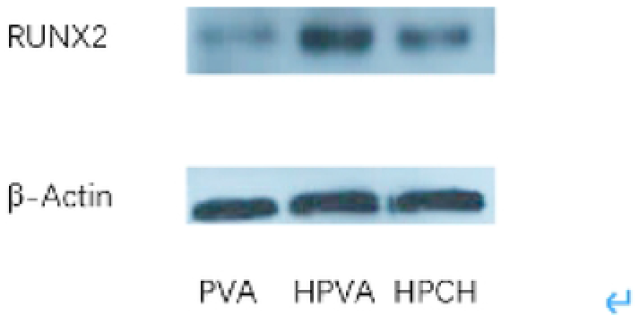
Western blotting of RUNX2

## 5. Conclusion

Bone scaffold materials are designed to elevate bone regeneration, scaffolds modification is applied to enhance cell adhesion, proliferation and differentiation. Surface characteristics of scaffolds hold pivotal role affecting its application. Surface porosity, area and roughness, as three features of scaffold surface, are usually chosen to be fabricated physically and chemically. The relationship between surface pore size and area is usually reverse, when the surface pore size increases, the corresponding surface area decreases. Manufacturing process can affect microporosity, then the surface area changed too. By adding different biomaterial particles, we can change the microporosity of scaffolds.

Scaffolds made of by electrospinning are widely used, the comprehensive discussions of the nanofiber scaffolds produced by electrospinning were conducted by former researchers (Xue et al.,2019). The advantages of adjustable fiber diameter, effective consumption of solution, high tensile strength, large surface area and porosity made electrospinning widely used in our daily life (Thenmozhi et al.,2017). such as Environmental applications, biomedical applications, applications in the field of chemistry and defence (Abdul et al.,2022). Biomedical applications of electrospinning nanofibers were developed to assist the wound healing, which is a deliberate process in tissue engineering including inflammation, proliferation, and remodeling phase(Nemati et al.,2019). Recently tissue engineering scaffolds from electrospinning attract a great area of interest and dramatic research. By incorporating various additives into nanofibers, we can modify the polymers then achieved special properties to work as extracellular matrix in tissue engineering (Dahlin et al.,2011;Victor et al.,2011).

Hydroxypropyl chitosan (HPCH) is an abundant and renewable resource found in most parts of the world, which makes it a cheap raw material for various applications(Ghannoum et al.,2015;Mani et al.,2005).

However, little research has been done on the use of HPCH and PVA as raw material within electrospinning. Blending of HPCH with PVA can not only change the microporosity and surface area of fabricated nanofibers, but also enhance the mechanical properties of the nanofiber mats. Characterization of the obtained nanofibers mats with respect to surface morphology along with the functional groups using SEM, FT-IR are studied(EI-Newehy et al.,2018). As shown in our study, the mechanical property and biocompatibility of the mixed nanofibers were enforced.

The composite results of this study indicate that the combined application of the HPCH and PVA can improve electrospun nanofiber characteristics, as demonstrated by the enhanced tensile strength and osteogenesis which contribute to increased microporosity and surface area. Taken together, these findings may translate into a more effective mats for bone regeneration. In conclusion, the results obtained in this study suggest that application of the HPCH and PVA can reinforce the nanofiber structure and thus induce osteoblast differentia.

